# CRISPR-Enabled Autonomous Transposable Element (CREATE) for RNA-based gene editing and delivery

**DOI:** 10.1101/2024.01.29.577809

**Authors:** Yuxiao Wang, Ruei-Zeng Lin, Meghan Harris, Bianca Lavayen, Bruce McCreedy, Robert Hofmeister, Daniel Getts

## Abstract

To address a wide range of genetic diseases, genome editing tools that can achieve targeted delivery of large genes without causing double-stand breaks (DSBs) or requiring DNA templates are necessary. Here, we introduce the CRISPR-Enabled Autonomous Transposable Element (CREATE), a genome editing system that combines the programmability and precision of CRISPR/Cas9 with the RNA-mediated gene insertion capabilities of the human LINE1 (L1) element. CREATE employs a modified L1 mRNA to carry a payload gene, and a Cas9 nickase to facilitate targeted editing by L1-mediated reverse transcription and integration without relying on DSBs or DNA templates. Using the system, a 1.1 kb gene cassette comprising an EF1α promoter and green fluorescent protein (GFP) gene is inserted into several genomic loci of multiple human cell lines. Mechanistic studies reveal that the CREATE system is highly specific with no observed off-target events. Together, these findings establish CREATE as a programmable gene delivery technology solely based on RNA components, enabling large-scale in vivo genome engineering with broad therapeutic potential.

## Introduction

The recent FDA approval of exagamglogene autotemcel (Casgevy®) for sickle cell disease has marked a significant milestone in CRISPR/Cas9-based gene therapies^1^. Traditional CRISPR/Cas9 systems, which introduce genomic double-strand breaks (DSBs), are best suited to disrupt defective genes and while CRISPR/Cas9 can also be used to insert new DNA templates via the homology-directed repair (HDR) pathway, the integration efficiency is low^2^. To avoid DSB-associated chromosomal deletions or translocations^3–6^, Base Editors (BEs) and Prime Editors (PEs), which perform gene edits by nicking a single DNA strand rather than creating DSBs, have been introduced^7,8^. BEs enable targeted single nucleotide conversions, while PEs can insert or replace up to 40 base pairs without a donor DNA template. Despite these improvements, these editors face limitations in payload capacity, which restricts their application in treating most genetic disorders involving a spectrum of mutations, insertions, or deletions across extensive genomic regions.

Expanding on the capabilities of PEs, twin-prime and template-jumping PEs have been developed to incorporate larger payloads of a few hundred base pairs (bps)^9^. Furthermore, combining PEs with sequence-specific serine recombinases has extended this capacity up to 36 kB for the delivery of large DNA fragments encoded in a donor plasmid^10,11^. Despite their potential, the technical complexity of these systems, which require the co-delivery of multiple components such as the PE editor with pegRNA, the serine recombinase, and a DNA donor plasmid, presents substantial challenges for clinical applications^12^.

The LINE1 retrotransposon (referred to as L1) is a mobile genetic element that comprises 17% of the human genome^13^. It naturally replicates via an RNA intermediate, capable of efficiently reverse-transcribing large mRNA sequences into cDNA and integrating into human genome^14,15^. L1 replication involves a bicistronic mRNA encoding two proteins, ORF1p and ORF2p^14^. The ORF1p is an RNA-binding protein that interacts with L1 mRNA transcripts^16^. ORF2p encodes an endonuclease (EN) and a reverse transcriptase (RT) domain^17,18^. A distinctive feature of L1 is its ‘cis preference,’ where ORF1p and ORF2p associate with their own mRNA transcript to form a ribonuclear protein complex (RNP)^19^, facilitating the reverse transcription and insertion of the L1 mRNA^20,21^. The endonuclease domain of ORF2p cleaves at a redundant 5’TTTT/AA3’ consensus sequence, initiating a target-primed reverse transcription (TPRT) process mediated by the RT domain of ORF2p that converts the L1 mRNA into cDNA and integrates into the genome^22,23^. The ORF2p RT domain was shown to be highly processive, capable of reverse transcription of long RNA sequences^23,24^. However, the consensus motif recognized by the EN domain of L1 ORF2p is found at numerous sites in the genome, precluding its application for targeted, programmable gene editing. A recent study attempting to combine Cas9 with another retroelement, the R2 retroelement via a direct fusion approach was unable to achieve complete retrotransposition and integration^25^.

Intrigued by the properties of L1, we developed the CRISPR-Enabled Autonomous Transposable Element (CREATE) system co-opting the transposable element for site-specific gene delivery. By combining the precision and programmability of CRISPR/Cas9 with the large-scale genome re-writing capability of L1, we inserted a 1.1 kb nucleotide sequence comprising a promoter and green fluorescent protein (GFP) into several sites in the human genome guided by Cas9. Together our findings show that the CREATE system is a programmable and highly specific gene delivery platform capable of delivering a payload of over 1 kb with only RNA components, thus offering considerable prospects for diverse therapeutic applications.

## Results

### Design of the CREATE editing system

The CREATE system encodes, on a single mRNA, the codon optimized L1 components ORF1p and ORF2p, separated by the native inter-ORF sequence which mediates an unconventional bicistronic expression of the two proteins^26^ (Fig. 1a). To facilitate nuclear import, the ORF2p is flanked with an N-terminal SV40 nuclear localization signal (NLS) and a C-terminal nucleoplasmin NLS, respectively. The inactivation of the catalytic residue within the EN domain of ORF2p (mutation D205A) is a critical modification to prevent DNA cleavage of the degenerate consensus motif recognized by natural L1. The payload cassette consisting of a promoter and the gene of interest is placed at the 3’UTR of the L1 mRNA to form the CREATE mRNA (Fig. 1a). To ensure that the payload cannot be expressed by direct translation prior to genome integration, the payload is encoded in the antisense orientation (Fig. 1a). Critically, the payload is flanked by sequences designed to hybridize with single-guide RNA (sgRNA) target sites in the genome to prime reverse transcription. These sequences are referred to as primer binding site 1 (PBS1) and reverse complement (RC) primer binding site 2 (RC-PBS2).

**Fig. 1.**
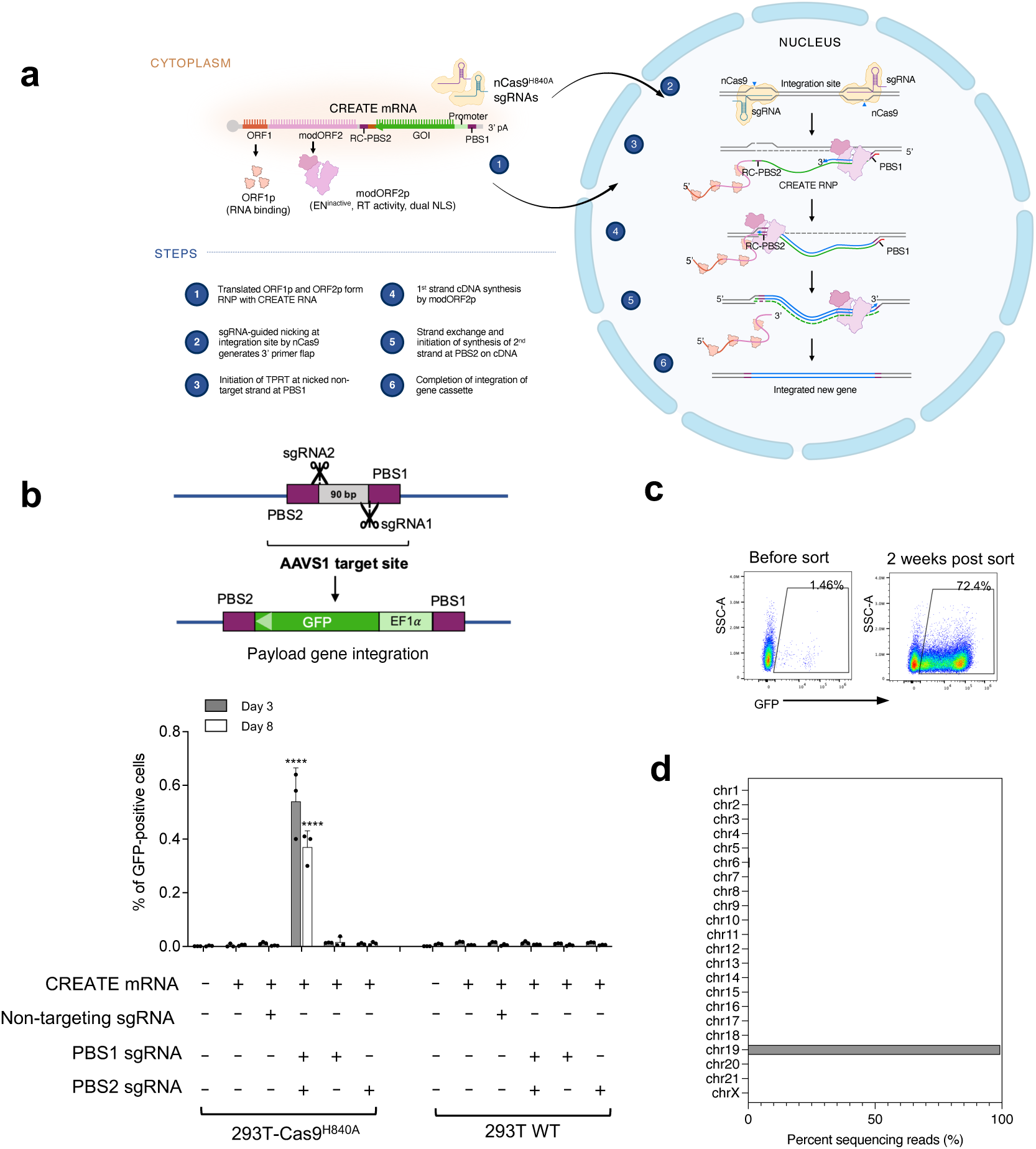
CREATE editing design and proof-of-concept in mammalian cells. **(a)** Design of the CREATE system and the mechanism for gene integration. (**b**) Proof-of-concept study demonstrating replacement of a 90 bp segment at AAVS1 locus with a GFP expression cassette using CREATE. Quantification of payload integration efficiencies shows successful editing requires Cas9 nickase activity and dual sgRNAs. Statistical analysis was performed using two-way ANOVA with Dunnett’s multiple comparisons test comparing against the first sample. **** indicates p < 0.0001. Statistical significance is labeled for samples with p <0.01. Data are mean ± SD. (**c**) AAVS1 edited cells before and after enrichment for GFP positive cells. (**d**) Genome-wide target hybridization sequencing confirms specific integration of GFP payload gene at AAVS1 locus on chromosomal 19. Identified potential sites with >5 sequencing reads were plotted across all chromosomes (see Supplementary Table 3 for details).

Upon cellular uptake, the CREATE mRNA is translated into ORF1p and ORF2p^ENdead^ proteins, which then co-assemble with the mRNA to form CREATE RNP and enter cell nucleus (Fig. 1a, step 1). Natural L1 retrotransposition relies on nicking of the non-targeting DNA strand by the EN domain of ORF2p^23,27^. To harness this mechanism, a Cas9^H840A^ non-target strand nickase is employed to introduce a single strand nick guided by sgRNA1 (Fig 1a, step 2). The liberated DNA 3’-flap then hybridizes with the PBS1 sequence on the CREATE mRNA, serving as the primer for the RT domain of ORF2p for first strand cDNA synthesis (Fig. 1a, step 3). The newly synthesized cDNA strand includes both the payload and the PBS2 sequences in the sense orientation (Fig. 1a, step 4). Subsequently, PBS2 hybridizes with the 3’ flap released from the second nicked site generated by sgRNA2, invoking the L1 template jump mechanism to complete the second strand synthesis and integration^23^ (Fig. 1a, step 5-6). The outcome of the editing cycle is the successful integration of the payload cassette between the PBS sites, discarding the L1 components, resulting in safe non-replicative gene insertion.

### CREATE-mediated large payload gene delivery in mammalian cells

To demonstrate that CREATE can be used to deliver functional genes, we designed a reporter payload cassette consisting of an EF1! core promoter-driven GFP with an SV40 poly(A) signal (totaling 1.1 kb). The AAVS1 site, a well-characterized genomic safe harbor site, was selected to show programmable targeted insertion. We chose two sgRNAs that have been validated previously, targeting sites within AAVS1 that are 90 bp apart^28^. Successful editing would replace the 90 bp DNA segment with the 1.1 kb payload cassette (Fig. 1b). Baldwin et al. showed that ORF2p can efficiently initiate TPRT with a 7-20 bp primer length^23^. Therefore, we designed PBS1 and RC-PBS2 in this range to anneal with the DNA flaps released from two sgRNA cut sites, and positioned them on either side of the payload.

Both Cas9 and ORF2p are large multidomain proteins, each exceeding 100 kDa. We were concerned that direct Cas9-ORF2p fusions would compromise activity and the ability to enter the nucleus. Since ORF2p could scan genomic DNA for entry sites, we hypothesized that it could independently identify sites nicked by Cas9, eliminating the need for a direct fusion. To test this concept, we developed a HEK293T cell line engineered to stably express a SpCas9 non-targeting strand nickase (293T-nCas9^H840A^). Fluorescence-activated cell sorting (FACS) was used to quantify the percentage of cells expressing GFP as a measurement of successful payload integration. Transfection of cells with CREATE mRNA and the two sgRNAs resulted in 0.5% and 0.4% GFP expression on Day 3 and Day 8, respectively (Fig. 1b). No GFP expression was observed in 293T-nCas9^H840A^ cells transfected with non-targeting control sgRNAs or in wildtype HEK293T cells (Fig. 1b). These results confirmed sgRNA-guided integration and ruled out the possibility of Cas9-independent, nonspecific insertion. Furthermore, we demonstrated that payload expression required co-transfection of both sgRNAs, as delivery of either sgRNA alone did not result in GFP positive cells (Fig. 1c). This result indicated that nicking on opposite DNA strands was necessary for TPRT and successful integration. Such requirement inherently enhances specificity, as a single independent sgRNA off-target binding would be insufficient to cause payload integration.

FACS based cell sorting was employed to enrich the CREATE edited cells to approximately 70% GFP positive, after which cells were maintained for an additional two weeks (Fig. 1c). During this time GFP expression remained stable (>70%), indicative of successful and stable genomic integration of the payload. To validate the site-specificity of the insertion and search for potential off-target editing, we performed next generation sequencing with target enrichment by using hybridization probes to capture EF1!-GFP payload sequences from genomic DNA extracted from edited cells. Insertion sites analysis across the whole genome showed highly specific editing at the intended AAVS1 locus at chromosome 19 (Fig. 1d). Detailed examination of the sequencing reads showed no off-target edits at other genomic regions (Fig. 1d and Supplementary Table 3). These results highlighted the target specificity of CREATE system.

To further appreciate the temporal requirements for delivery of the CREATE components, sequential transfection of RNA for the individual CREATE elements was performed. The specific protocols included (1) Simultaneous transfection of CREATE mRNA and sgRNAs (One-shot Protocol); (2) Transfecting CREATE mRNA four hours prior to sgRNAs (Two-step, mRNA-first Protocol); (3) Transfecting sgRNAs four hours before CREATE mRNA (Two-step, sgRNA-first Protocol). In Protocols (2) and (3), culture media was replaced immediately prior to the second transfection. Observations supported superior performance of the two-step protocols when compared to the one-shot approach (Supplementary Fig. 1a).

Notably, the mRNA-first strategy yielded the highest CREATE editing efficiency, reaching ∼1.2% GFP expression on day 3 and remaining stable on day 8. The improvement is likely due to the necessary time for the ORF1p and ORF2p mRNA to be translated, co-assemble with CREATE mRNA and subsequently enter the cell nucleus. Delivery of sgRNAs before the CREATE RNP complex forms might result in the single strand nicks being repaired by the cell before the CREATE editing process can occur. A time interval of 4 to 8 hr between mRNA and sgRNA transfections yielded the highest editing rates of approximately 1.5%. Likewise, an increase of sgRNA above 0.242 μg per each well improved the editing efficiency (Supplementary Fig. 1b and 1c).

### CREATE editing is predicated on L1 retrotransposition mechanism

Mutational analysis of the key components of CREATE was performed to investigate the proposed editing mechanisms. We first examined whether CREATE editing required non-target strand (Cas9^H840A^) rather than target strand nickase (Cas9^D10A^) activity. When we transfected CREATE mRNA and the two AAVS1-targeting sgRNAs into HEK293T cells stably expressing either Cas9^D10A^ or Cas9^H840A^ nickase, we observed GFP expression and integration exclusively in the presence of Cas9^H840A^ nickase (Fig. 2a), confirming the natural L1 replication mechanism that starts with non-target strand cleavage. Of note, the activity of Cas9^H840A^ also destroys the PAM sequence after successful insertion, ensuring a single allele may only be edited once, mitigating concerns about the potential for continuous nicking of the edited genome. Next, we introduced a D702A mutation to inactivate the RT domain of L1 ORF2p (CREATE^RTdead^). This modification entirely abolished the integration and expression of the GFP payload (Fig. 2a) confirming L1 RT activity was required for cDNA synthesis. Similarly, the removal of the entire EN domain (CREATE mRNA^ΔEN^) resulted in a marked decrease in editing efficiency, indicating that while the EN domain must be rendered inactive to avert cleavage of the genome at redundant consensus sites recognized by L1 naturally, its complete removal was detrimental to the editing process. This is consistent with the recent reports showing that the EN domain and RT domain of L1 ORF2p are highly integrated structurally and functionally^23,27^.

**Fig. 2.**
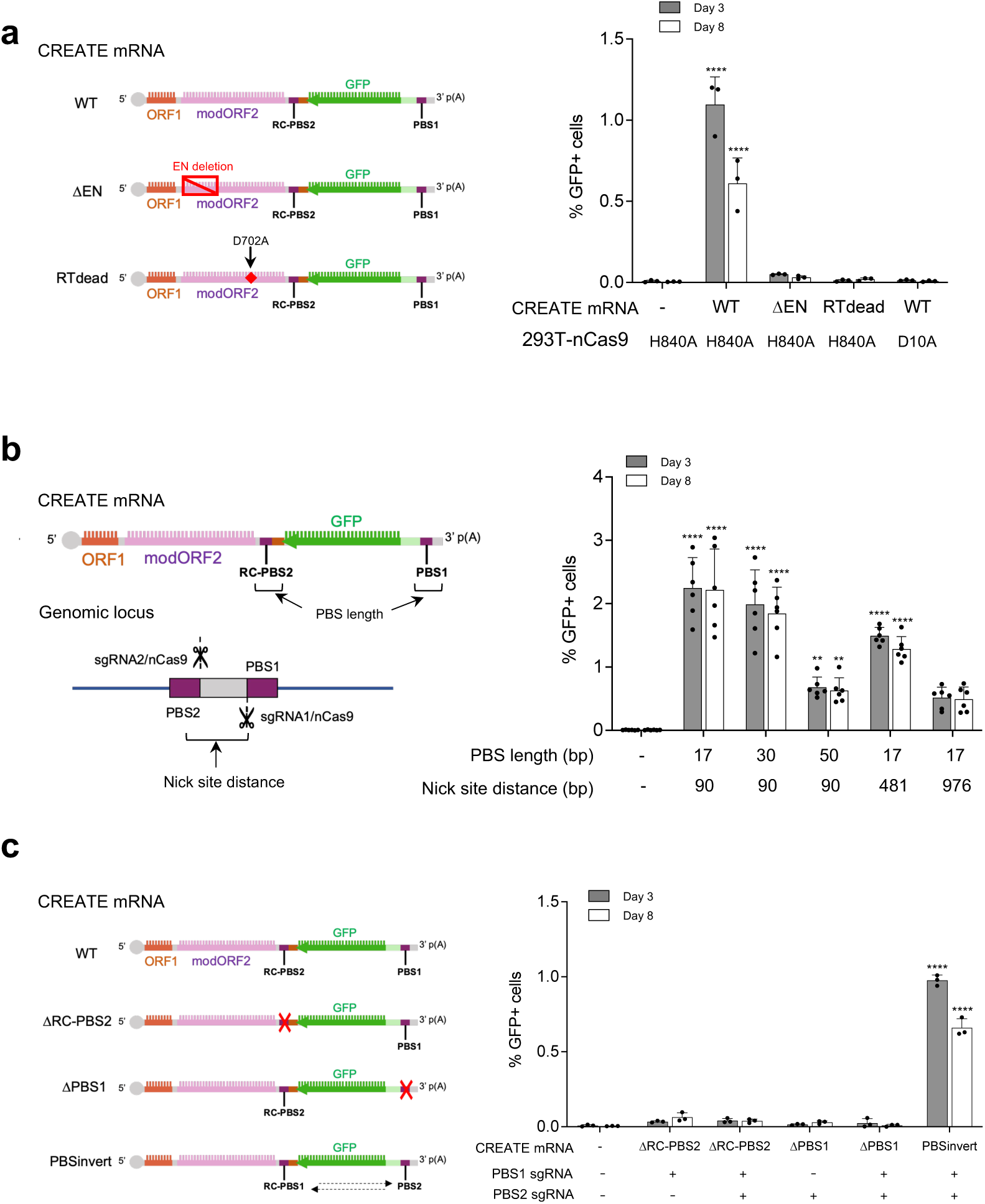
Mechanistic analysis of CREATE editing. (**a**) Mutational analysis of the key enzymatic activities and protein domains involved in editing. (**b)** Influence of the length of PBS sequences in CREATE mRNA and the distance between sgRNA nick sites in the genome on editing efficiency. (**c**) CREATE editing requires two PBS sequences that matches the two sgRNA target sites. Statistical analysis was performed using two-way ANOVA with Dunnett’s multiple comparisons test comparing against the first sample. **** p < 0.0001.** p<0.01. Statistical significance is labeled for samples with p <0.01. Data shown are representative of at least two independent experiments. All data are mean ± SD.

### sgRNA target sites and PBS sequences influence editing efficiency

To determine if PBS length impacts insertion efficiency, we designed a series of CREATE mRNA constructs with different PBS1 and RC-PBS2 hybridization sequence lengths (17 bp, 30 bp, and 50 bp) and measured insertion efficiency at the AAVS1 sites targeted with the same sgRNAs. In these experiments 17-30 bp appeared to support the most efficient editing, while 50 bp was associated with a reduction of GFP integration efficiency (Fig. 2b). These results are with a recent study investigating the poly-A length required for efficient TPRT by L1 ORF2^27^.

As a search-and-replace tool, CREATE swaps out the genomic sequence between sgRNA1 and sgRNA2 for the payload (Fig. 1b). Next, we investigated the impact of the distance between the two sgRNA nicking sites on editing efficiency. Three sgRNA1s with binding sites located 90 bp, 481 bp or 976 bp from a fixed site targeted by sgRNA2 were designed. While CREATE achieved successful integration of a GFP cassette and replacement of the segment between the sgRNAs in all cases, the editing efficiency dropped slightly with increasing distance between the sgRNAs (Fig. 2b).

### A requirement for paired sgRNAs and matched PBS sequences impart high fidelity and precision on CREATE

To demonstrate the importance of two PBS sites flanking the payload, we designed CREATE mRNA with either PBS1 or a RC-PBS2 sequence. In the presence of AAVS1 targeting sgRNAs, the removal of PBS1 or RC-PBS2 resulted in a lack GFP expression on day 3 or day 8 (Fig. 2c). Additionally, exchanging the position of PBS1 and RC-PBS2 while simultaneously encoding them in a reverse-complementary sequence (“RC-PBS1” and “PBS2) restored the GFP expression. The requirement for the two PBS sequences that matched the two sgRNA cut sites were critical for a highly specific integration. When we assessed the top 5 predicted off-target sites for sgRNA1 and sgRNA2 by PCR, no integration of GFP cassette into theses potential sites was observed (Supplementary Fig. 2). Together, these data validate the proposed mechanism of CREATE and show that the editing process has several inherent checkpoints that prevent off-target activity. These include: 1) correct base-pairing of PBS1 with sgRNA1 cut sites; 2) correct base-pairing of reverse transcribed RC-PBS2 with sgRNA2 cut site and 3) adjacent sgRNA1 and sgRNA2 nicking.

### CREATE-mediated gene integration into different genomic loci

To determine the ability of the CREATE system to edit and deliver genes specifically into sites in the genome beyond AAVS1, we engineered CREATE to deliver GFP to other genomic loci including HEK3, PRNP, and IDS. The HEK3 locus, situated on chromosome 1 is frequently chosen to benchmark gene editing tools^29–31^. The PRNP locus on chromosome 20 encodes the prion protein implicated in multiple neurodegenerative diseases^32,33^, and IDS gene on chromosome X is associated with Hunter’s syndrome^34^, a lysosomal storage disease. For each targeted site, a pair of sgRNAs 70-90 bp apart were selected. CREATE mRNA, which carried the corresponding 30 bp PBS1 and RC-PBS2 flanking the GFP expression cassette, was deployed to each target site. Flow cytometric analysis on day 3 demonstrated all three sites exhibited comparable editing efficiency, as determined by GFP expression (Fig. 3a).

**Fig. 3.**
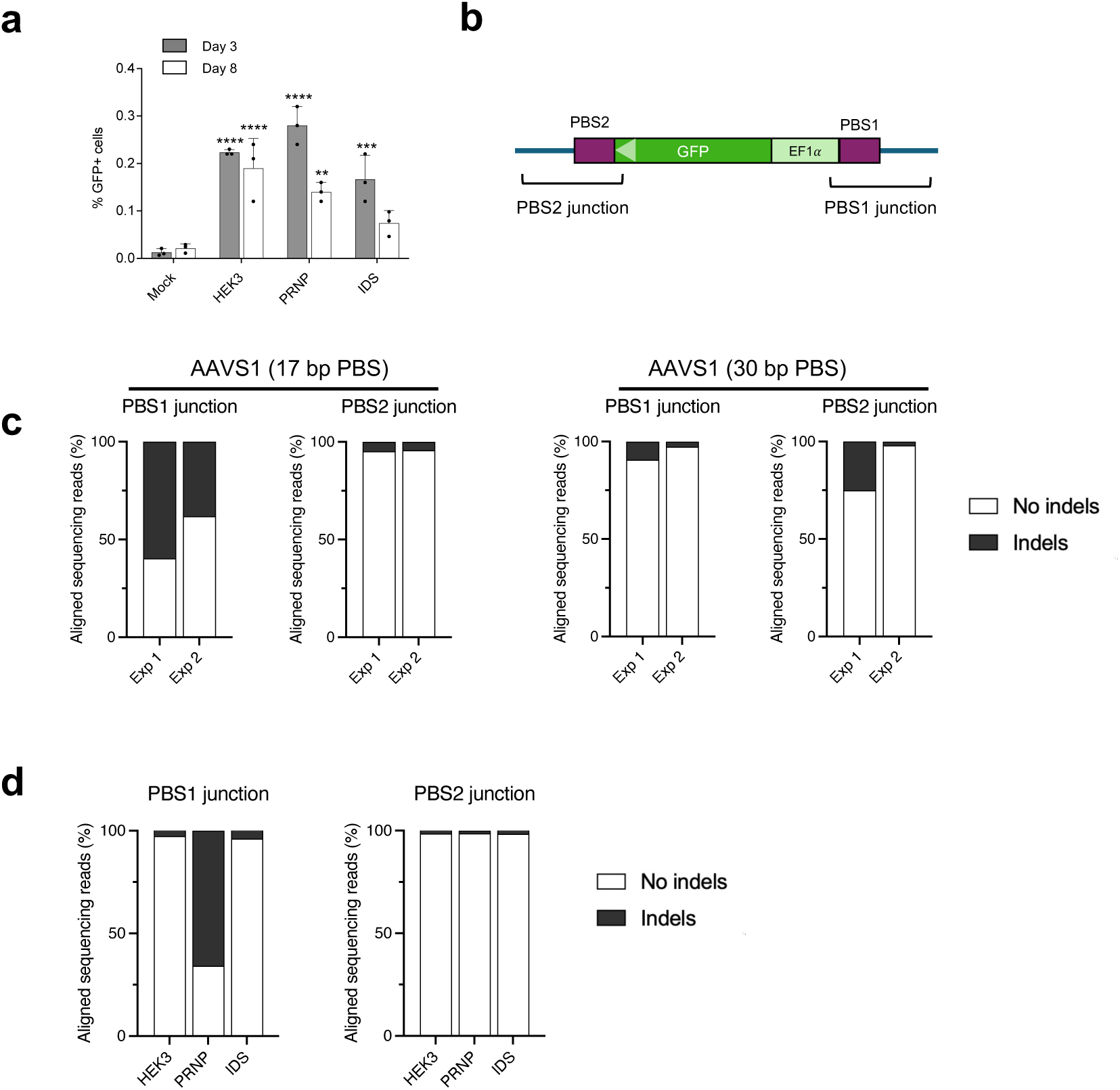
CREATE as a fully programmable gene delivery tool in mammalian cells. (**a**) Editing of three additional genomic loci by designing unique sgRNAs and matching PBSs. Statistical analysis was performed using two-way ANOVA with Dunnett’s multiple comparisons test comparing against the first sample. **** p < 0.0001. *** p<0.001.** p<0.01. Statistical significance is labeled for samples with p <0.01. Data shown are representative of at least two independent experiments. All data are plotted as mean ± SD. (**b**) Amplicon sequencing to characterize the junction regions of integrated payload genes. (**c**) The percentage of indels at the integration junction at AAVS1 locus. Two different CREATE mRNAs were designed with 17 bp or 30 bp PBSs, both targeting AAVS1 locus. Exp 1 and Exp 2 represent two independent experiments. (**d**) The percentage of indels at integration junctions at HEK3, PRNP and IDS loci. Edited cells were from experiment shown in Fig. 3a. Data from (**c**) and (**d**) were analyzed and plotted using CRISPResso2^48^.

To characterize the editing accuracy at the junction sites between payload sequences and genomic sequence, we performed amplicon sequencing by designing PCR primers to cover the expected junctions (Fig. 3b). Interestingly, for AAVS1 edited cells we observed varying levels of indel formation at the PBS1 junction, while PBS2 junction was >95% without indels (Fig. 3c). Changing of the length of PBS from 17 bp to 30 bp reduced indel formation at PBS1 (Fig. 3c). We also performed the same analysis on the junction regions for cells with targeted insertion at HEK3, PRNP and IDS sites. The results showed that for HEK3 and IDS, the insertion were more than 95% accurate; while the PRNP site showed indel formation only at the PBS1 junction but not PBS2 (Fig. 3d). Taken together, these results showed that the formation of indels occurs only at the junction site and are likely dependent on the sgRNA sequence for the corresponding target locus. Optimization of target sgRNAs for each particular target site is necessary to reduce junction infidelity and improve editing efficiency.

### CREATE as a fully RNA-based gene delivery tool for different mammalian cell types

Finally, an important attribute of CREATE gene editing is the ability to deliver all components as RNA into mammalian cells (Fig. 4a). To demonstrate this, we synthesized mRNA encoding nCas9^H840A^ with N- and C-terminal NLS. Electroporation was used to deliver the nCas9 mRNA, CREATE mRNA and two sgRNAs designed to target AAVS1 locus into HEK293T cell and the immortalized liver cell line Huh7. We found that one-time electroporation led to ∼1% and 0.2% GFP payload re-expression in Huh7 and HEK293T cells, respectively (Fig. 4b). Taken together, these results highlight the potential to develop CREATE into a fully RNA based gene delivery tool for in vitro and in vivo applications.

**Fig. 4.**
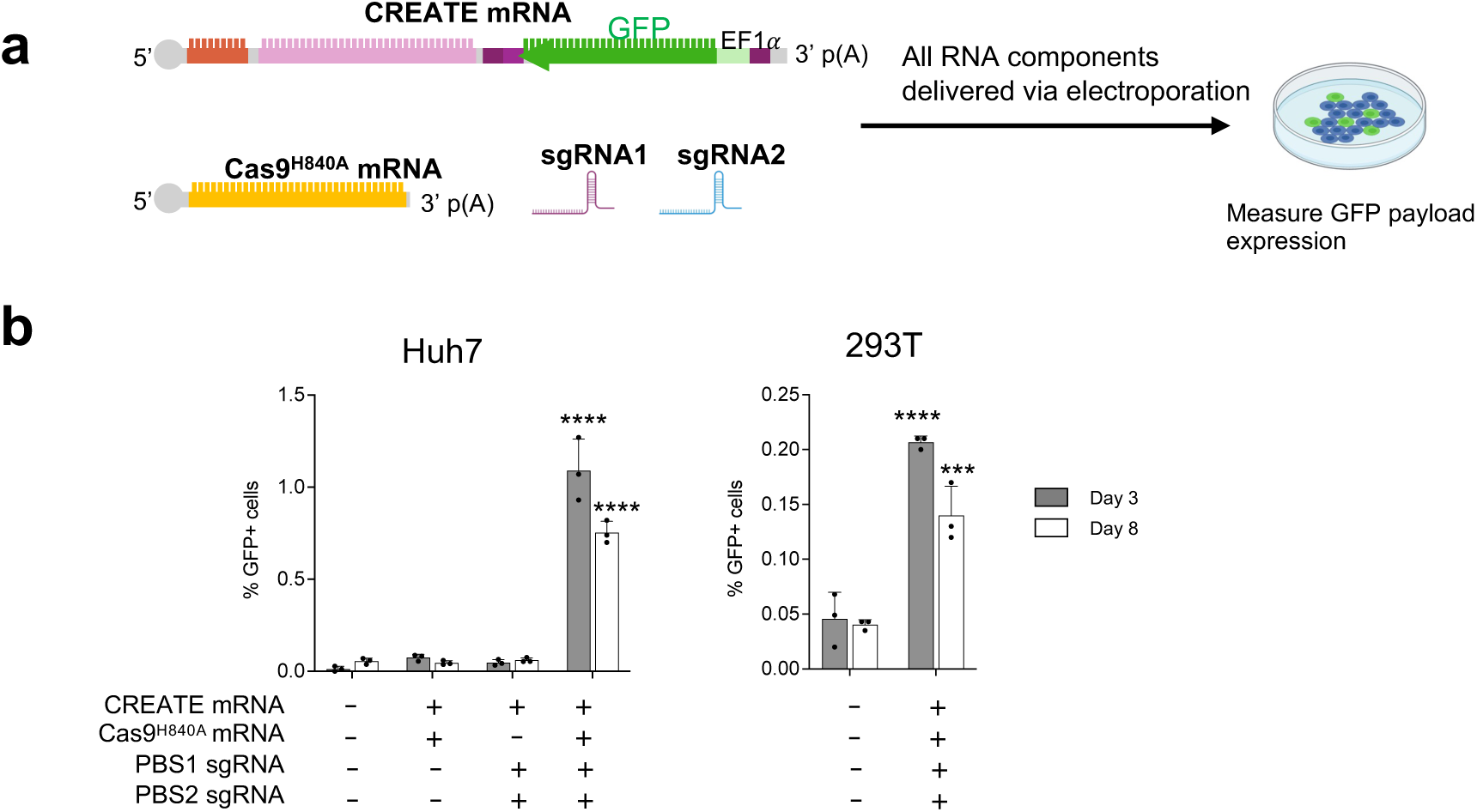
CREATE as a fully RNA-based gene delivery system to multiple cell types. (**a**) Co-delivery of CREATE editing components as mRNA/sgRNA. (**b**) RNA-based GFP payload insertion at AAVS1 locus in Huh7 and HEK293T cells. Statistical analysis was performed using two-way ANOVA with Dunnett’s multiple comparisons test comparing against the first sample. **** p < 0.0001. *** p<0.001. Statistical significance is labeled for samples with p <0.01. Data shown are representative of at least two independent experiments. All data are plotted as mean ± SD.

## Discussion

Here we have engineered CREATE, a programmable, site-specific tool for integration of full-length genes into the human genome, based on the human L1 retrotransposon. Key design elements for the site specificity of CREATE include the modification of the natural L1 ORF2p endonuclease and the introduction of a Cas9 nickase. Guided by two sgRNAs, Cas9 nickase targets the desired integration sites without DSBs, thereby mitigating the risk of chromosomal instabilities. Two primer binding sites flanking the payload are critical for the initiation of reverse transcription and converting the payload RNA into cDNA for integration into the genome. This is the first evidence of using L1 or any other retrotransposon system to deliver genes with specificity and programmability. An earlier study explored the direct fusion of a Cas9 or Cas12a endonuclease with the reverse transcriptase domain of L1 ORF2p for site-specific integration^35^. While Cas-directed cleavage and subsequent RNA payload integration was achieved, the study was limited to payload insertion into high-copy plasmids in *E. coli*. The use of an active Cas9 endonuclease exhibited significant toxicity. In contrast, CREATE employs a Cas9 nickase in combination with the full-length, endonuclease inactivated ORF2p for reverse transcription. Recent high-resolution cryo-EM structures of ORF2p complexed with DNA and RNA substrates shed light on the process of target site recognition and initiation of reverse transcription^22,23^. Multiple domains in ORF2p fold into an integrated structure that wrap around the substrates to effectively couple genomic DNA binding, target site cleavage and target-primed reverse transcription. Our findings support the notion that preserving all essential domains of ORF2p can facilitate productive integration in mammalian genomes.

Non-LTR retroelements have gained increasing interest for the delivery of larger payloads. R2 retroelements, found in metazoans and birds, have been used to insert large RNA payloads (up to 4kb) into eukaryotic genomes. However, R2 elements are naturally restricted to the ribosomal DNA (rDNA) loci, which limits their utility for editing in other genomic locations. In addition, insertion at rDNA site may result in reduced expression over time due to rDNA array instability^36^. An attempt to integrate R2 elements with Cas9 for RNA-guided reverse transcription and integration was unsuccessful to achieve complete cDNA synthesis and integration^25^. Exploiting the natural replication mechanisms of an engineered L1 element, CREATE achieved an integration efficiency of approximately 1.5% in multiple mammalian cell lines, inserting a 1.1 kb GFP cassette without detectable off-target integration in multiple genomic sites. Given L1’s natural capacity to mediate insertion of approximately 6 kb of RNA, we anticipate that future iterations of CREATE could deliver even larger sequences.

The L1 element has been associated with increased risk of cancer and cellular senescence^37,38^. A gradual loss of epigenetic suppression of L1 transcription, can lead to reactivation of L1 retrotransposition which can cause insertional mutagenesis or chronic inflammation triggered a type I interferon response^39–41^. In the present study, no off-target integrations were observed suggesting that CREATE does not cause genomic instability. Several properties and modification ensure that the CREATE system integrates the payload gene without propagating natural L1 components or perturbing their repressed state. CREATE mRNA is only transiently expressed and once on-target integration has occurred, the sgRNA binding sites are altered which prevents any further re-editing event. Finally, the EN domain of ORF2p in the CREATE system is inactivated to prevent non-specific cleavage of genomic DNA.

Varying degrees of indel formation at the transgene/genomic locus junction were noted. This phenomenon has also been described for PE-related technologies that utilize double-nicking. 10-20% indel by-products were observed for the PE3 system^8^. GRAND editing, a PE-derived technology, yielded 5.8% to 63.0% accurate editing with substantial indel formation^42^. Systematic optimization of pegRNA design and manipulation of cellular DNA repair pathways can reduce indel by-products^43,44^. We anticipate that a more in-depth understanding of the CREATE integration mechanism along with sgRNA and target site optimization will enhance junction fidelity and overall editing efficiency. Furthermore, the combination of the modified L1 element with alternative programmable RNA-guided DNA endonucleases such as Cas12, TnpB and Fanzor can be explored to expand the capability of the CREATE system^45–47^

In summary, the development of CREATE combines the gene integration capacity of a human L1 retroelement with the CRISPR/Cas9 precision. As a fully RNA-based genome engineering tool that does not require DNA templates, it complements existing approaches and has the potential to address a large variety of genetic diseases.

## Methods

### Cell culture

HEK293T (ATCC, cat. no. CRL-3216) and Huh7 cells (JBRC, Cat. No. JCRB0403) were maintained in Dulbecco’s Modified Eagle’s Medium (DMEM) supplemented with 10% (v/v) fetal bovine serum (FBS, Gibco) and 1% (v/v) Penicillin/Streptomycin/Glutamine (Gibco). Cells were cultured at 37°C with 5% CO_2_. TrypLE Select (Thermo Fisher Scientific) was used for passaging and harvesting cells.

### sgRNAs

sgRNAs were acquired from Synthego or Thermo Fisher Scientific and reconstituted with TE buffer into 100 pmol/µL stock solution. Sequences of sgRNAs are listed in Supplementary Table 1.

### Generation of Cas9-expressing HEK293T cell lines

PiggyBac vectors expressing Cas9 nickase H840A or D10A variants with an N-terminal SV40 NLS and a C-terminal Nucleoplasmin NLS under the CMV promoter were constructed by Vector Builder. PiggyBac vectors (1.25 µg) and hyPBase mRNA (0.7 µg) were co-transfected into 2.5×10^6^ HEK293T cells using MaxCyte ATx electroporator by following the manufacturer’s protocol. Three days after transfection, cells were selected with hygromycin (400 µg/mL) for 10 days, to acquire stable Cas9-expressing HEK293T cell lines.

### Cloning of CREATE constructs

DNA sequences encoding human L1 ORF1p (GenBank: AAB59367.1) and ORF2p (GenBank: AAA51622.1) with N-terminal SV40 NLS and C-terminal nucleoplasmin NLS were codon optimized and synthesized at Twist Bio. Endogenous L1 interORF region sequences flanked by restriction sites PshAI (GACAGCCGTC) and AccI (GTATAC) was synthesized at Twist Bio. The full constructs containing all L1 components including ORF1p, interORF and ORF2p were then assembled using Hifi Assembly mix (NEB) into pCMV vector with the CMV promoter region removed. A T7 promoter was introduced upstream of ORF1p for in vitro transcription. The payload cassette including an EF1! core promoter, EGFP and SV40 poly(A) signal was synthesized by Integrated DNA Technologies and inserted at the 3’UTR of the L1 mRNA using Hifi Assembly mix (NEB). gBlocks gene fragments containing different PBS site sequences were synthesized by Integrated DNA Technologies, and subsequently cloned into the vector to flank the payload gene. Sequences of PBS sites and payloads are listed in Supplementary Table 2. Mutation of ORF2 RT domain was introduced using Q5 site-directed mutagenesis kit (NEB). Deletion of ORF2 EN domain was performed by PCR and ligation cloning.

### mRNA synthesis via in vitro transcription

Plasmid templates for *in vitro* transcription were linearized by restriction enzyme and templates were purified by column clean up (GeneJET PCR Purification Kit, Thermofisher) and eluted in 40µl of nuclease free water. All in house generated mRNA was transcribed via T7 polymerase and co-transcriptionally capped utilizing New England Biolabs HiScribe T7 mRNA Kit with CleanCap Reagent AG according to the manufacturer’s protocols. The in vitro transcription reaction was allowed to proceed for 2 hours at 37 °C. Reaction mixtures were treated with DNase I and incubated at 37 °C for 30 minutes before immediate purification (Monarch® RNA Cleanup Kit, NEB). A sample of each mRNA was reserved for quality analysis and transcripts were tailed with *E. coli* poly(A) polymerase (NEB) at 37 °C for 30 minutes, and subsequently purified (Monarch® RNA Cleanup Kit, NEB). All tailed and untailed mRNA samples were normalized to a concentration of 50ng/µl and quality was assessed via TapeStation RNA ScreenTape Analysis (Agilent, RNA ScreenTape, 5067-5576) according to the manufacturer’s protocol.

### Lipofectamine transfection

HEK293T cells were seeded on 24-well plates at 80,000 cells the day prior to transfection in DMEM with 2% FBS. Transfection was performed by either one of these protocol:

- One-shot Protocol (#1): L1-mRNA (3 ug per well) and sgRNAs (0.121 ug each sgRNA per well) were transfected using Lipofectamine MessengerMAX (Invitrogen).
- Two-step, mRNA first Protocol (#2): L1-mRNA (3 ug per well) was transfected using Lipofectamine MessengerMAX. Transfected cells were incubated at 37°C with 5% CO_2_ for 4 hour. After refreshing the media, cells were then transfected with sgRNAs (0.121 ug each sgRNA per well) using Lipofectamine RNAiMax (Invitrogen).
- Two-step, sgRNA first Protocol (#3): sgRNAs were transfected using Lipofectamine RNAiMax. Transfected cells were incubated at 37°C with 5% CO_2_ for 4 hour. After refreshing the media, cells were then transfected with L1-mRNA (3 ug per well) using Lipofectamine MessengerMAX.

### Electroporation

Huh7 cells were harvested by TrypLE Select and resuspended in MaxCyte buffer at a density of 6.25 x 10^7^ cells/mL. For MaxCyte electroporation, 15 µg of L1-mRNA, 7.5 µg of nCas9 mRNA, and 2.5 µg/each dual sgRNAs were gently mixed with 1.25 x 10^6^ Huh7 cells in a total volume of 27.5 µL MaxCyte buffer. 25 µL of cell-RNA mixture was transferred into an OC-25×3 cuvette. Electroporation was carried out on the MaxCyte ATX at the indicated electroporation energy. Immediately after electroporation, the OC-25×3 cuvettes with cells were incubated at 37°C for 15 min to allow for cell membrane recovery. Finally, cells were added to a 6-well plate feeded with DMEM with 5% FBS for the first 3 days after electroporation. The medium was removed and switched back to DMEM with 10% FBS after day 3. Cells were harvested on day 3 and day 8 post-transfection for FACS analysis.

### Flow cytometry analysis and cell sorting

Flow cytometry analysis was performed on day 3 and day 8 after transfection. Transfected cells were collected after PBS washing and TrypLE Select digestion and resuspended in PBS with 1% FBS and 2mM EDTA for flow cytometry analysis (CYTEK). Data were analyzed by FlowJo 10.9.0 software.

For experiments that require enriched GFP-positive cells, cells were collected three to five days after transfection, and sorted by flow cytometry (BD FACSMelody sorter) to obtain the GFP-positive cells. Sorted cells were cultured in DMEM with 20% FBS for the initial 7 days to facilitate cell recovery.

### Genomic DNA extraction

One million collected cells were centrifuged at *5,000 g* for 5 min at 4°C and the pellet was resuspended in 100-µL cold PBS. Genomic DNA was extracted using Monarch Genomic DNA Purification Kit (NEB) according to manufacturer’s instructions and eluted in 20 µL water.

### Target hybridization sequencing

Genomic DNA (gDNA) extracted was used to prepare Insertion Site Analysis (ISA) libraries using the Twist Target Enrichment procedure. Briefly, gDNA was enzymatically fragmented, followed by end repair, dA-tailing and adapter ligation, and library amplification using indexed primers. The resulting pre-hybridization libraries underwent QC followed by hybridization-capture with baits corresponding to the transgene/insert, and a final library amplification using universal primers. The final library was sequenced by Illumina platform using 2×150bp sequencing configuration in paired end mode.

For data analysis, raw BCL files generated by the sequencer were converted to raw fastq files for each sample using bcl2fastq v.2.20. Fastq reads were then trimmed with adapter sequences. Homology analysis was performed for vector genome against human reference genome (build hg38) and high homology regions identified in reference human genome were hard masked. A combined reference genome consisting of vector genome and hard masked human reference was generated and reads were aligned to the combined genome using bwa v0.7.4. The bam file was subsequently processed with Samtools v0.1.16. Finally, integration sites were identified by taking into account pairs of reads that support insertions and alignment to both vector and human reference genomes. Identified potential insertion sites were clustered and annotated by Azenta proprietary ISA pipeline. Potential insertion sites with >5 confirmed reads per location were analyzed and plotted. Detailed analysis of identified insertions sites with >5 confirmed reads are shown in Supplementary Table 3.

### Amplicon sequencing

To validate the site-specific integration of CREATE payload, target regions were PCR amplified and analyzed by sequencing methods. Genomic DNA samples were PCR amplified with Q5 High-Fidelity 2X PCR Master Mix (NEB) based on the manufacturer’s protocol. PCR amplicon primers were designed specifically to integration locus-CREATE junctions and are listed in Supplementary Table 4. Amplicons were analyzed by 1% agarose gel electrophoresis and purified according to the sizes using the DNA Gel Extraction Kit (NEB).

To quantify integration of CREATE payload by targeted deep sequencing, amplicons from PCR amplification of genomic-CREATE insertion junctions were prepared for sequencing on a MiSeq (Illumina). Alignment of amplicon reads to a reference sequence was performed using CRISPResso2. Reads with a mean quality score <30 were discarded.

### Determination of CREATE editing activity at potential off-target sites

To evaluate the off-target activity of CREATE editing, the potential off-target sites of a single sgRNA were detected by targeted PCR amplification. Potentially top-ranking predicted off-target sites of spacers were selected using the CRISPR-Cas9 guide RNA design checker (Integrated DNA Technologies). PCR primers and predicted off-target insertion are listed in Supplementary Table 5. The PCR reactions were performed using Q5 High-Fidelity 2X PCR Master Mix (NEB) with optimized Tm and the PCR products were detected by 1% agarose gel.

### Statistics and reproducibility

At least three replicates were analyzed in each independent experiment to ensure the experimental data were reliable. Data are expressed as mean ± s.d. Statistical analyses were performed with GraphPad Prism (version 9.0, GraphPad Software).

## Data availability

Sequences including primers, sgRNAs are available in the supplementary materials.

## Acknowledgments

We thank R. Vale (HHMI Janelia) & S. Mukherjee (Columbia U) for critical reading of the manuscript and the entire Myeloid Therapeutics team for support.

## Author contributions

Conceptualization: YW, DG, RL, BM

Methodology: YW, DG, RL, MH

Experiments and Data Analysis: RL, MH, BL, ND, YW

Writing: YW, RL, DG, RH

