## Supplementary information for "CRISPR-Enabled Autonomous Transposable Element (CREATE) for RNA-based gene editing and delivery"

### Supplementary Materials

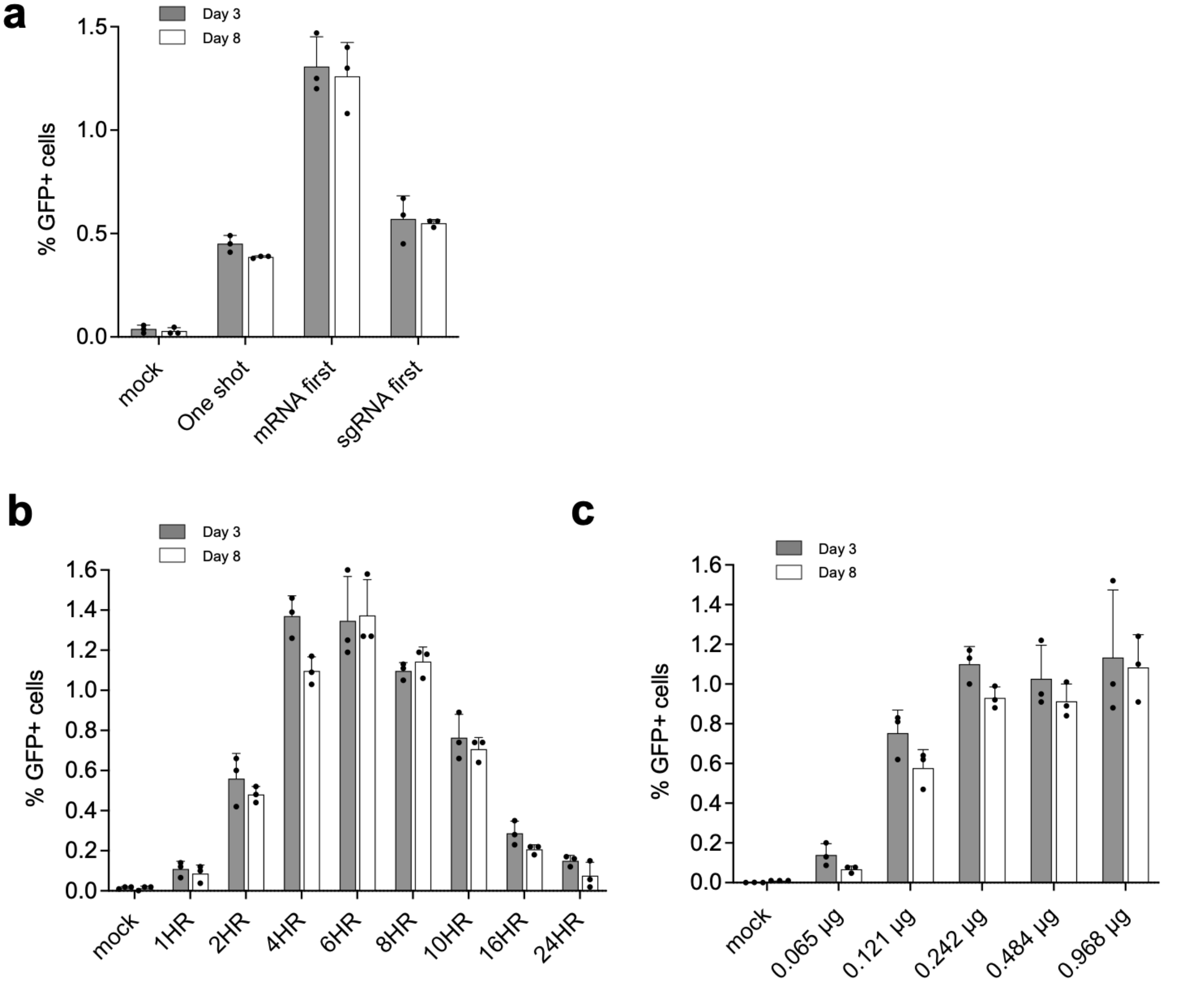


**Fig. S1. Optimization of transfection protocol to improve CREATE editing efficiency.** (a) Optimization of transfection protocol improved editing efficiency. Protocol 1 (one shot), Protocol 2 (mRNA first) and Protocol 3 (sgRNA first). Data shown are representative of at least two independent experiments. (b) Optimization of the incubation time between mRNA and sgRNA transfections. (c) Optimization of the total amount of CREATE sgRNAs transfected.

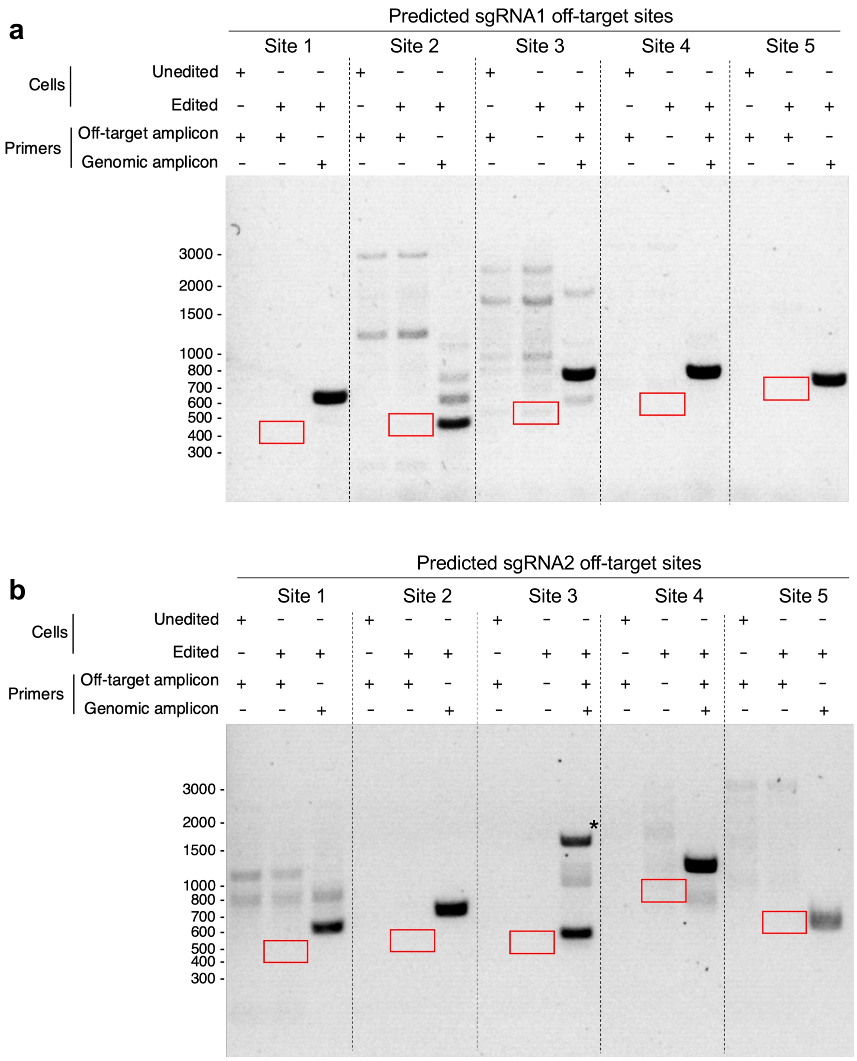


**Fig. S2. PCR detection of potential off-target editing in AAVS1 loci edited cells.** Top 5 predicted off-target sites for sgRNA1 and sgRNA2 were examined. Red box indicate the size of the expected PCR product if off-target integration occurred. * indicate non-specific band amplicon.

**Supplementary Table 1. Sequences of sgRNAs used in this study.**

| **Nicking sgRNA** | **Spacer sequence (5'-3')** |
|---|---|
| AAVS1_nicking_PBS1 | GAUGGAGCCAGAGAGGAUCC |
| AAVS1_nicking_PBS2 | GCAGCUCAGGUUCUGGGAGA |
| AAVS1_nicking_PBS1_481-bp replacement | GCUCUUCCAGCCCCCUGUCA |
| AAVS1_nicking_PBS1_976-bp replacement | CCUUCCCUGCCGCCUCCUUC |
| HEK3_nicking_PBS1 | GUCAACCAGUAUCCCGGUGC |
| HEK3_nicking_PBS2 | GGCCCAGACUGAGCACGUGA |
| PRNP_nicking_PBS1 | GCAGUGGUGGGGGGCCUUGG |
| PRNP_nicking_PBS2 | GCAUGUUUUCACGAUAGUAA |
| IDS_nicking_PBS1 | GCAUUUUCGAUUCCGUGACU |
| IDS_nicking_PBS2 | ACUGAGGGAUGUCUGAAGGC |
| Negative Control, non-targeting | AAAUGUGAGAUCAGAGUAAU |

#### Supplementary Table 2. Sequences of PBS sites and payloads used in this study.

| **EF1$\alpha$-GFP payload in sense direction (1083 bp)** |
|---|
| EF1$\alpha$ core promoter-GFP-SV40 polyA signal<br>GGGCAGAGCGCACATCGCCCACAGTCCCCGAGAAGTTGGGGGGAGGGGTCGGCAATTGATCCGGTGCCTAGAGAAGGTGG<br>CGCGGGGTAAACTGGGAAAGTGATGTCGTGTACTGGCTCCGCCTTTTTCCCGAGGGTGGGGGAGAACCGTATATAAGTGCAG<br>TAGTCGCCGTGAACGTTCTTTTTCGCAACGGGTTTGCCGCCAGAACACAGGGTTTAGTGAACCGTCAGATCCCGCCACCATG<br>GTGAGCAAGGGCGAGGAGCTGTTCACCGGGGTGGTGCCCATCCTGGTCGAGCTGGACGGCGACGTAAACGGCCACAAGTT<br>CAGCGTGTCCGGCGAGGGCGAGGGCGATGCCACCTACGGCAAGCTGACCCTGAAGTTCATCTGCACCACCGGCAAGCTGCC<br>CGTGCCCTGGCCCACCCTCGTGACCACCCTGACCTACGGCGTGCAGTGCTTCAGCCGCTACCCCGACCACATGAAGCAGCAC<br>GACTTCTTCAAGTCCGCCATGCCCGAAGGCTACGTCCAGGAGCGCACCATCTTCTTCAAGGACGACGGCAACTACAAGACCC<br>GCGCCGAGGTGAAGTTCGAGGGCGACACCCTGGTGAACCGCATCGAGCTGAAGGGCATCGACTTCAAGGAGGACGGCAAC<br>ATCCTGGGGCACAAGCTGGAGTACAACTACAACAGCCACAACGTCTATATCATGGCCGACAAGCAGAAGAACGGCATCAAGG<br>TGAACTTCAAGATCCGCCACAACATCGAGGACGGCAGCGTGCAGCTCGCCGACCACTACCAGCAGAACACCCCCATCGGCGA<br>CGGCCCCGTGCTGCTGCCCGACAACCACTACCTGAGCACCCAGTCCGCCCTGAGCAAAGACCCCAACGAGAAGCGCGATCAC<br>ATGGTCCTGCTGGAGTTCGTGACCGCCGCCGGGATCACTCTCGGCATGGACGAGCTGTACAAGTAAAACTTGTTTATTGCAGC<br>TTATAATGGTTACAAATAAAGCAATAGCATCACAAATTTCACAAATAAAGCATTTTTTTCACTGCATTCTAGTTGTGGTTTGTCCA<br>AACTCATCAATGTATCTTA |

| **CREATE constructs** | **PBS1 sequence (5'-3')** | **RC-PBS2 sequence (5'-3')** |
|---|---|---|
| AAVS1_90-bp replacement_17-bp PBS | TCCTCTCTGGCTC | GCAGCTCAGGTTCTGGG |
| AAVS1_90-bp replacement_30-bp PBS | TCCTCTCTGGCTCCATCGTAAG CAAACCTT | CAGCCGCGTCAGAGCAGCTCAG GTTCTGGG |
| AAVS1_90-bp replacement_50-bp PBS | TCCTCTCTGGCTCCATCGTA AGCAAACCTTAGAGGTTCT GGCAAGGAGAG | GATCAGTGAAACGCACCAGACA GCCGCGTCAGAGCAGCTCAGGT TCTGGG |
| AAVS1_90-bp replacement_17-bp PBS reverse | CCCAGAACCTGAGCTGC | GAGCCAGAGAGGA |
| AAVS1_481-bp replacement_17-bp PBS | CAGGGGGCTGGAAGAGC | GCAGCTCAGGTTCTGGG |
| AAVS1_976-bp replacement_17-bp PBS | TCCTCTCTGGCTC | GCAGCTCAGGTTCTGGG |
| HEK3_90-bp replacement_30-bp PBS | CGTGCTCAGTCTGGGCCCCAA GGATTGACC | CCCAGCCAAACTTGTCAACCAG TATCCCGG |
| PRNP_72-bp replacement_30-bp PBS | AGGCCCCCCACCACTGCCCCA GCTGCTGCA | TTGGGGTAACGGTGCATGTTTTC ACGATAG |
| IDS_70-bp replacement_30-bp PBS | CACGGAATCGAAAATGCTTCA GAAGGTTCT | TTGTCAGAATTCCACTGAGGGAT GTCTGAA |

**Supplementary Table 3. Target hybridization sequencing results for AAVS1 edited samples**

| Chromosome | Location | Strand | Total_Reads | Annotation |
|---|---|---|---|---|
| chr17 | 38905912 | + | 7 | Endogenous Human EF1$\alpha$ sequence |
| chr19 | 55115486 | - | 26904 | On-target insertion at AAVS1 (chr19: 5515280 – 5515880) |
| chr19 | 55115589 | + | 15692 | |
| chr19 | 55115800 | - | 132 | |
| chr19 | 55115148 | - | 8 | |
| chr19 | 55115690 | - | 8 | |
| chr19 | 55115285 | - | 5 | |
| chr2 | 32916556 | - | 21 | Mis-alignment to repetitive sequence |
| chr2 | 32916333 | - | 8 | Mis-alignment to repetitive sequence |
| chr5 | 14653390 | + | 8 | Endogenous Human EF1$\alpha$ pseudo gene 13 sequence |
| chr6 | 73521212 | + | 134 | Endogenous Human EF1$\alpha$ sequence |
| chr6 | 73521001 | - | 117 | |

*Identified potential insertion with >5 reads at a specific location are analyzed*

**Supplementary Table 4. Sequences of primers used for genomic DNA junction amplification.**

| Target | Forward primer (5'-3') | Reverse primer (5'-3') |
|---|---|---|
| AAVS1-PBS1 | CGGAGCCAGTACACGACATC | TCCCAGGGCCGGTTAATGTG |
| AAVS1-PBS2 | GGGCTGGCTACTGGCCTTAT | ATCACTCTCGGCATGGACGA |
| PRNP-PBS1 | CGGAGCCAGTACACGACATC | CTGGAGGCAACCGCTACC |
| PRNP-PBS2 | GTGGTTGTGGTGACCGTGT | ATCACTCTCGGCATGGACGA |
| HEK3-PBS1 | CGGAGCCAGTACACGACATC | TGATGTGGGCTGCCTAGAAA |
| HEK3-PBS2 | GCCCTGAGATCTTTTCCTCTGT | CAACGAGAAGCGCGATCACA |
| IDS-PBS1 | AAAGAACGTTCACGGCGACT | CTGTTTCAGGCAGGCAATCC |
| IDS-PBS2 | GTTGGCAAAACTCAAGGCATCA | ACATGGTCCTGCTGGAGTTCG |

#### Supplementary Table 4. Sequences of primers used for PCR to detect off-target insertion.

| Name | Target sequence | Location | GFP insertion forward primer (5'-3') | Genomic reverse primer (5'-3') | Genomic forward primer (5'-3') | Genomic amplicon size (bp)[#] | Predicted off-target amplicon (bp)[##] |
|---|---|---|---|---|---|---|---|
| AAVS1_nicking_PBS1 | GATGGAGCCAGAGAGGATCC | chr19:-55115573 | n/a | n/a | n/a | n/a | n/a |
| PBS1_off_target_1 | GATGAAGTCAGAGAGGATCC | chr20:-2451191 | GCTTGCCGTAGGTGGCAT | CACTCTGGCGTTAAAGGAGCA | GAAGTCACCCCCGAATCAGC | 649 | 427 |
| PBS1_off_target_2 | CACGGAGCCAGGGAGGATCC | chr9:-127595186 | GCTTGCCGTAGGTGGCAT | CAGGATCTTACCCTGCCATGA | AGACTCGGAGCTCAAACTGC | 439 | 453 |
| PBS1_off_target_3 | GCTGGAGGCAAGAGAGGATCC | chr11:-75540340 | GCTTGCCGTAGGTGGCAT | CATCACAATGCCCCAGGACT | GTCCCTTCACAGACCATGCG | 757 | 513 |
| PBS1_off_target_4 | GATGAATGCTGAGAGGATCC | chr1:-93567284 | GCTTGCCGTAGGTGGCAT | GCCGCTTGTGACGTTAATGG | ACCTCCTCATCTTGGCACATC | 779 | 529 |
| PBS1_off_target_5 | GTGGGAGGCAGAGAGAATCC | chr1:-11579676 | GCTTGCCGTAGGTGGCAT | GGCATTCCCTGTGAGAGTGT | GCCCTCTGTCAATAACCGCT | 741 | 667 |
| AAVS1_nicking_PBS2 | GCAGCTCAGGTTCTGGGAGA | chr19:+55115469 | n/a | n/a | n/a | n/a | n/a |
| PBS2_off_target_1 | GAGGCTCCGGTTCTGGGAGA | chr1:+198494971 | CGGGATCACTCTCGGCATGG | GCAGAGCTACCGGCTCCATG | CCTTTCACTGTGAACCTCATTGTTAGC | 559 | 460 |
| PBS2_off_target_2 | TCAGTTCAGGTTCTGGAAGA | chr10:+6483525 | GCATGGACGAGCTGTACAAGTAAAA | GCCCTCGGGGTCCCTATTTA | CCTTGATGTGGCATTATTAACAGTGC | 662 | 556 |
| PBS2_off_target_3 | GCTGGCTCTGGTTCTGGGAGA | chr11:+132450011 | GCATGGACGAGCTGTACAAGTAAAA | GCACCAACAGGTTCAGAGGT | GTGCTTTGGAGGACGACCAG | 429 | 521 |
| PBS2_off_target_4 | GCCCCTCGGGTCCTGGGAGA | chr19:+44516904 | GCATGGACGAGCTGTACAAGTAAAA | CGATTCCCCATATAGCTCACTCC | GGTGCATCACGTTTGGCTTT | 953 | 809 |
| PBS2_off_target_5 | GCATTTCTGTTTCTGGGAGA | chr2:+30765077 | CGGGATCACTCTCGGCATGG | GGCACGTGACCATCCCTTAAT | GGAACGCTTCCAATTCACCC | 567 | 577 |

[#] Genomic amplicon size: If no off-target insertion happens, the PCR amplification using corresponding Genomic forward primer and Genomic reverse primer will produce PCR band of indicated size.
[##] Predicted off-target amplicon size: If off-target insertion happens based on the prediction, the PCR amplification using GFP insertion forward primer and Genomic reverse primer will produce PCR band of indicated size.
